# TLR7 and RIG-I dual-adjuvant loaded nanoparticles drive broadened and synergistic responses in dendritic cells *in vitro* and generate unique cellular immune responses in influenza vaccination

**DOI:** 10.1101/2020.07.17.207423

**Authors:** Randall Toy, M. Cole Keenum, Pallab Pradhan, Katelynn Phang, Patrick Chen, Chinwendu Chukwu, Anh Nguyen, Jiaying Liu, Sambhav Jain, Gabrielle Kozlowski, Justin Hosten, Mehul S. Suthar, Krishnendu Roy

## Abstract

Although the existing flu vaccines elicit strong antigen-specific antibody responses, they fail to provide effective, long term protection – partly due to the absence of robust cellular memory immunity. We hypothesized that co-administration of combination adjuvants, mirroring the flu-virus related innate signaling pathways, could elicit strong cellular immunity. Here, we show that the small molecule adjuvant R848 and the RNA adjuvant PUUC, targeting endosomal TLR7s and cytoplasmic RLRs respectively, when delivered together in polymer nanoparticles (NP), elicits a broadened immune responses in mouse bone marrow-derived dendritic cells (mBMDCs) and a synergistic response in both mouse and human plasmacytoid dendritic cells (pDCs). In mBMDCs, NP-R848-PUUC induced both NF-κB and interferon signaling. Interferon responses to co-delivered R848 and PUUC were additive in human peripheral blood mononuclear cells (PBMCs) and synergistic in human FLT3-differentiated mBMDCs and CAL-1 pDCs. Vaccination with NPs loaded with H1N1 Flu antigen, R848, and PUUC increased percentage of CD8+ T-cells in the lungs, percentage of antigen-specific CD4+T-cells in the spleen, and enhanced overall cytokine-secreting T cell percentages upon antigen restimulation. Also in the spleen, T lymphopenia, especially after in vitro restimulation, was observed. Our results demonstrate that simultaneous engagement of TLR7 and RIG-I pathways using particulate carriers is a potential approach to improve cellular immunity in flu vaccination.

**GRAPHICAL ABSTRACT:** 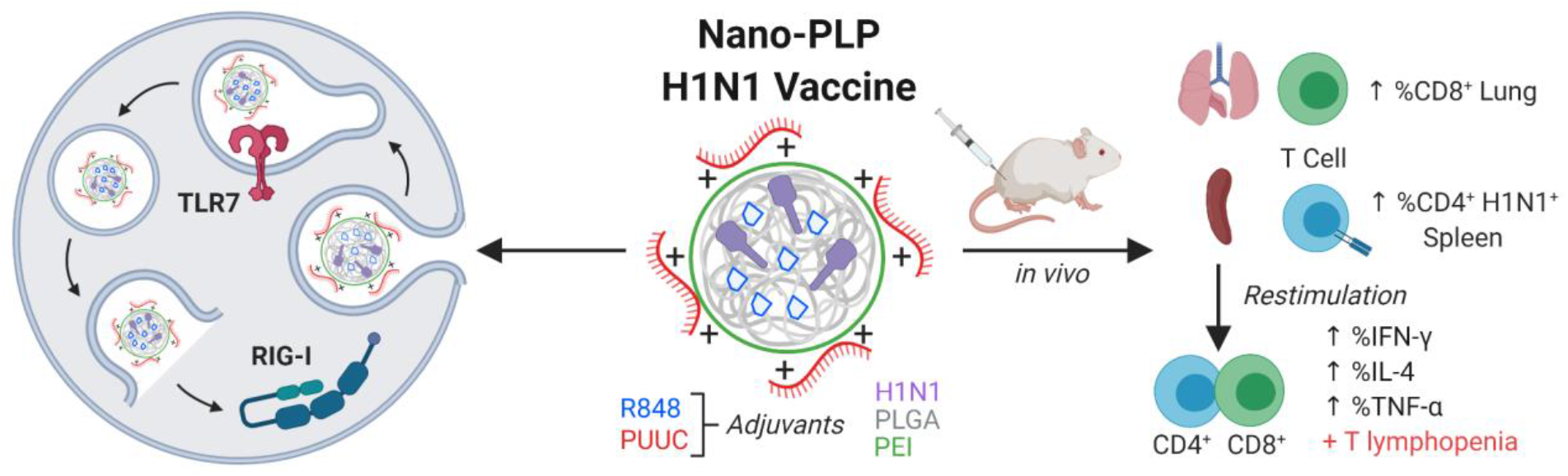

## INTRODUCTION

It is estimated that every year, globally, between 294,000 and 518,000 people die of influenza and associate complications. In the previous century, three global pandemic outbreaks of influenza occurred, with the largest killing an estimated 50-100 million.^1–3^ While vaccines have controlled the spread of other deadly diseases, such as smallpox, polio, and measles, vaccines against the flu, although somewhat effective in the short-term, do not offer long-term protection. In the elderly, it is estimated that flu vaccines only protect 46% of patients against the 2009 pandemic H1N1 virus.^4^ High affinity antibodies generated in response to flu vaccines primarily recognize the surface hemagglutinin (HA), which mutates frequently. As a result, it is difficult to achieve long-lasting neutralizing antibody responses.^5^ Designing vaccines that can elicit robust induction of persistent T cell immunity may enable better protection against the flu.^6,7^

Virus-specific CD8+ and CD4+ T cells are more likely to induce memory formation to effectively combat a wider flu virus repertoire. To boost the efficacy of flu vaccines, non-specific immune adjuvants have been deployed to enhance antibody and cell-mediated responses.^8^ Antibody-specific responses to the A(H1N1)pdm09 antigen were significantly boosted by an α-tocopherol oil-in-water emulsion-based adjuvant in both young and middle-aged adults at 21 days and 42 days, but not after 12 months.^9^ Children have benefited from flu vaccines co-delivered with the MF-59 adjuvant, which strengthens the antibody response.^10^ In the absence of a universal influenza vaccine, immune adjuvantation represents a major strategy to improving flu vaccinations.

Second generation immune adjuvants that target specific innate immune receptors are powerful weapons for combating cancer and infectious disease.^11,12^ These adjuvants carry pathogen-associated molecular patterns (PAMPs), which trigger Toll-like receptors (TLRs) and cytosolic immune-receptors (e.g. retinoic-inducible gene 1 (RIG-I) like receptors (RLRs), cyclic GMP-AMP Synthase (cGAS)). For specific application to flu vaccines, combinations of various synthetic TLR-agonists have been reported, but combinations of TLR agonists with RLR or cGAS-ligands have not been thoroughly investigated.^13–15^ RLRs play a critical role in antiviral defense against the flu, West Nile virus, and Zika virus.^16–20^ RLRs recognize RNA sequences with 5’ triphosphorylated (5’ppp) ends, which is a hallmark of viral RNA. RLR activation induces a conformational change that allows caspase activation and recruitment domains (CARDs) to interact with the adaptor protein MAVS, which drives a signaling cascade that initiates Type I interferon production. Split vaccines adjuvanted with IRF3-activating small molecule RLR agonists have improved antibody titers and lethal challenge survival in mouse models of influenza.^21^ Activation of RIG-I may also enhance cell-mediated immunity by improving the T cell priming capability of pDCs.^22^ RIG-I ligands have been co-administered with HA antigens, which resulted in enhanced germinal center reactions and follicular helper T cell response kinetics at sparing doses.^23^ When combined with virus-like particles expressing hemagglutinin (HA) and neuraminidase antigens from the H5N1 virus, RIG-I ligands enhanced Th1 cytokine levels in CD4+ T cells and the antibody response induced by the virus-like particles.^24^

Since combination adjuvants have been shown to improve immune responses, we hypothesized that concurrent activation of endosomal TLR7 and the cytoplasmic innate immune sensor RIG-I using polymeric particulate carriers would initiate broadened or synergistic innate immune responses that enhance cellular immunity against flu. We defined a “broadened response” as when a combination of adjuvants induces a larger number of types of immune responses than a single adjuvant (e.g., IL-6 and IFN-β production compared to only IL-6 production). A “synergistic response” is defined as when the response to the combination of adjuvants is greater than the sum of the responses to two adjuvants delivered individually (e.g. a combination of two adjuvants triggers production of 3 ng ΙFN-β when each adjuvant alone only triggers production of 1 ng IFN-β). In contrast, an additive response to combination adjuvants would be precisely the sum of the responses to two individual adjuvants. This adjuvant combination is relevant to the physiological makeup of the flu virus, which is a ssRNA that activates TLR7 and RLRs in the absence of non-structural proteins inhibiting innate immunity.^25,26^ Specifically, we hypothesized that the concurrent activation of TLR7, which triggers NF-κB mediated pro-inflammatory pathways, and RIG-I, which triggers interferon signaling pathways critical to anti-viral immunity, would generate an innate immune activation profile that improves vaccine responses more effectively than either adjuvant by itself. Our initial goal was to develop an efficient system that delivers both adjuvants to dendritic cells, which are a critical bridge between innate and adaptive immunity.^27^ We developed a pathogen-like nanoparticle (PLP) system that could co-deliver antigens and multiple adjuvants on a single particle, can be internalized efficiently by murine bone marrow-derived dendritic cells (mBMDCs), escape the endolysosomal pathway, and induce Type I interferon production through delivery of a RIG-I adjuvant (PUUC) to activate cytosolic RLRs. We then demonstrated that PUUC could broaden the immune response to the TLR7 adjuvant R848 in mBMDCs. In addition, we found that R848 induces synergistic interferon responses in pDCs. The R848 and PUUC combination induced broadened and synergistic antibody and cell-mediated responses when delivered on nanoparticles with encapsulated HA antigen to mice. This broadened and synergistic response, according to previous adjuvant studies, could result in longer lasting protection.^22^ Taken together, our results indicate that simultaneous engagement of the TLR and RLR pathways could be beneficial for anti-viral cellular immunity and the development of more potent, next-generation vaccines.

## MATERIALS AND METHODS

### Murine bone marrow-derived dendritic cell (mBMDC) culture

Bone marrow was harvested from the tibias and fibulas of BALB/c or C57 Bl/6 mice (6-10 weeks, Jackson Labs, Bar Harbor, ME). For RIG-I^−/−^ mBMDCs, knockout mice of a mixed C57Bl/6J and 129×1/SvJ background were kindly provided by the lab of Michael Gale Jr. at the University of Washington, Seattle.^28^ The bone marrow cells were processed through a 40 μm cell strainer, treated with RBC lysis buffer, and seeded in Petri dishes at a concentration of 1,000,000 cells/mL. GM-CSF differentiated mBMDCs were cultured by growing murine bone marrow-derived cells in RPMI media (Invitrogen) with 10% characterized fetal bovine serum (HyClone, Logan< UT), 1% penicillin-streptomycin, 2 mM glutamine, 1x beta-mercapethanol, 1 mM sodium pyruvate, and 20 ng/mL murine recombinant GM-CSF (Peprotech, Rocky Hill, NJ). Media was refreshed on days 2, 4, and 6. Experiments were performed with cells in culture for 7 days. Human FLT-3 differentiated mBMDCs were cultured by growing murine bone marrow-derived cells with 10% characterized fetal bovine serum (HyClone, Logan< UT), 1% penicillin-streptomycin, 2 mM glutamine, 1x beta-mercapethanol, 1 mM pyruvate, and 200 ng/mL human recombinant FLT-3 (Peprotech, Rocky Hill, NJ) for 9 days. Experiments were performed immediately with human FLT-3 differentiated murine BMDCs.

### Microparticle and nanoparticle synthesis and characterization

Hydrophobic TLR7 adjuvant (R848) was purchased from STEMCell Technologies (Vancouver, CA). The RIG-I adjuvant PUUC (MW = 60,200 g/mole) was synthesized in house as described in a published protocol.^29^ Luciferase mRNA (MW = 130,000 g/mole) was synthesized by the lab of Philip Santangelo at the Georgia Institute of Technology. Fluorescent labeling of the mRNA for uptake and endosomal escape studies was achieved by a previously published method.^30^ PLGA nanoparticles were synthesized using a double emulsion method as described in a published protocol.^31^ Resomer^®^ RG 502 H Poly(D-Lactide-Co-Glycolide) (Sigma-Aldrich, St. Louis, MO) was dissolved in dichloromethane and water in a 1:20:5 w/v/v ratio to form a primary emulsion, which was sonicated at 65% power for 2 minutes. The secondary emulsion was formed by adding the primary emulsion to a solution of 5% polyvinyl alcohol (87-89% hydrolyzed, Sigma Aldrich) in a 5:16 v/v ratio, which was sonicated at 65% power for 5 minutes. To remove dichloromethane by evaporation, the formulation was stirred in a fume hood for 3 hours. Large PLGA particles were removed by centrifugation for 20 minutes at 3500 g. PLGA nanoparticles were pelleted by ultracentrifugation at 80,000 g for 20 minutes and then washed once with DI H2O. The nanoparticles were then formed by coating the PLGA nanoparticles with branched polyethylenimine (Polysciences, MW=70,000, Warrington, PA) through reaction with EDC and sulfo-NHS as described in previous work.^31^ The nanoparticles were washed once with 1 M NaCl solution and once with DI H2O by ultracentrifugation for 20 minutes. Next, the nanoparticles were sonicated in a bath sonicator for 10 minutes prior to lyophilization in RNAse free DI H2O for 48 hours. PUUC loading was achieved by pipetting of a solution of PUUC adjuvant in RNAse free DI H2O into a suspension of nanoparticles in sodium phosphate buffer (pH 6.5) treated with diethyl pyrocarbonate (DEPC). After mixing, the nanoparticles and adjuvant were vortexed for 1 minute. To complete the loading process, the nanoparticles with adjuvant were rotated at 4° C overnight. In dose escalation studies, the mass of nanoparticles was held constant for all PUUC doses. Measurement of nanoparticle size before PEI modification and zeta potential after PEI modification were measured using a Malvern Zetasizer. R848 loading was measured by dissolving nanoparticles in DMSO followed by absorbance measurements at 324 nm on a BIOTEK plate reader. Loading of PUUC was quantified using the Nucleic Acid Quantification module of the Gen5 software on a BIOTEK Plate Reader.

### Endosomal escape assay

Glass coverslips were sterilized and placed into 24 well plates. Murine BMDCs were seeded on the coverslips at a density of 100,000 cells/well and allowed to grow overnight. Fluorescently labeled nanoparticles were added to the cells and incubated for 2 hours. Cells were washed 3x with PBS and media was replaced. The cells were returned to the incubator for 0, 2, or 24 hours. At each timepoint, cells were washed 3x with warmed PBS and fixed with BD Cytofix buffer for 10 minutes. Cells were then washed 3x with warmed PBS and permeabilized with 0.002% Triton X for 10 minutes. After 3 more washes with PBS, the cells were blocked with 10% donkey serum overnight at 4 C. Following overnight incubation, cells were washed 3x with PBS and incubated with a cocktail of primary antibodies to intracellular compartment markers - clathrin (1:500, Biolegend MMS-423P), caveolin (1:200, Santa Cruz sc-53564), CD63 (1:50, Abcam ab193349), EEA1 (1:100, Santa Cruz sc-365652), and LAMP1 (1:200, Abcam ab25630) – for 30 minutes at 37 C. Cells were washed 3x with PBS and then incubated with an Alexa-Fluor 488 donkey-anti-mouse secondary antibody (1:250, Thermo Fisher) for 30 minutes at 37 C. Next, cells were washed 3x with PBS and the coverslips were mounted onto glass slides using Prolong Gold with DAPI. Slides were imaged on a Perkin Elmer Spinning Disk microscope with an EM-CCD camera (60x magnification). Manders M1 overlap coefficients (PLGA-PEI nanoparticles to intracellular compartments) were determined using the Co-localization toolbox in Volocity software for individual cells.

### *In vitro* activation of murine BMDCs with adjuvant-loaded nanoparticles

On day 7 of culture, GM-CSF-derived mBMDCs were plated at a density of 300,000 cells/well in 96-well plates and allowed to settle for 2 hours before the addition of nanoparticles with adjuvant. After treatment with nanoparticles, supernatants were harvested at 24 hours. ELISA or Luminex was used to measure cytokine concentrations after cell activation (IFN-β, IL-1β, TNF-α, CXCL10). FLT-3-derived mBMDCs were analogously plated and treated with nanoparticles at day 9 of culture. Cell culture supernatant was harvested at 24 hours and IFN-β was measured by ELISA.

### *In vitro* activation of CAL-1 pDC with adjuvant-loaded nanoparticles

CAL-1 pDCs were provided as a gift from Dr. Takahiro Maeda of Nagasaki University and Dr. Dennis Klinman of NIH. The cells were cultured in RPMI medium, 0.3µg/mL L-glutamine, 25 mM HEPES, 10% heat-inactivated characterized fetal bovine serum, and 1% penicillin/streptomycin. Cells were plated at a density of 150,000 cells/well in a 96 well plate. After treatment with nanoparticles, supernatants were harvested at 24 hours. Luminex was used to measure human cytokine concentrations after cell activation (IFN-α, IFN-β, and TNF-α).

### *In vivo* adaptive immune response to adjuvant-loaded nanoparticles

BALB/c mice (12 weeks old) were used to measure the adaptive immune response. Mice were immunized by intramuscular injection (50 μL in each quadricep muscle, for a total 100 μL injection volume) with PLGA-PEI nanoparticles loaded with H1N1 flu antigen (Influenza A H1N1 (A/California/04/2009)/HA0 protein(full length), Sino Biological, China), H1N1 flu antigen + R848, H1N1 flu antigen + PUUC, or H1N1 flu antigen + R848 + PUUC on days 0 and 28. The mice were sacrificed on day 35 and lungs, blood, popliteal lymph nodes, and spleens were collected. Serum was isolated from blood by centrifugation at 10,000 g for 5 minutes at 4 C. Antibody titers (IgG1, IgG2a) were measured in serum with ELISA. Lymph nodes were passed through a 40 μm strainer to make single cell suspensions and were stained for the B220 marker. Lungs and spleens were homogenized with a GentleMACS dissociator and passed through a 40 µm strainer. Lung cells were stained for CD3 and CD8 and analyzed by flow cytometry. Splenocytes were either stained for flow cytometry or seeded on 96-well plates for restimulation assays. The Class II I-A(d) Influenza HA2 96-104 AELLVLLEN tetramer was obtained from the NIH Tetramer Core Facility (Emory Vaccine Center, Atlanta, GA). Class II tetramer-specific CD8-(presumed CD4+) T-cell, CD4+ central memory T-cell (CD44+ CD62L+ CCR7+), CD4+ effector memory T-cell (CD44+ CD62L-CCR7-), and CD8+ central memory T-cell (CD62L+ CD127+ KLRG+) populations were measured with flow cytometry. For restimulation, cells were incubated with flu antigen at a concentration of 50 ng/mL for 6 hours while incubated with brefeldin. Cells were stained for CD3, CD4, CD8, IFN-α, IL-4, and TNF-α and were analyzed by flow cytometry.

### Statistical analysis

All statistical analyses were performed using GraphPad Prism 8. Datasets were analyzed for normality using the Shapiro-Wilk normality test. To assess the statistical significance of the difference between two normal datasets, a Mann-Whitney test was performed. To assess the statistical significance of the difference between three or more normal datasets, a one-way ANOVA was performed, and multiple comparisons were evaluated using Tukey’s test.

## RESULTS

### Nano-PLPs, but not micro-PLPs, efficiently deliver the RIG-I adjuvant PUUC into the cytosol of mouse BMDCs and induce Type I interferon production

To generate PLPs, we synthesized PLGA microparticles (MPs) and nanoparticles (NPs) for the delivery of encapsulated R848 adjuvant and surface-loaded RIG-I RNA adjuvant (PUUC). Surface loading of the negatively charged PUUC adjuvant was facilitated by conjugation of the cationic polymer branched polyethylenimine (PEI) to the particle surface (**Figure 1A, Table 1**). Before PEI modification and subsequent PUUC loading, PLGA MPs were approximately 2.4 μm in diameter, while PLGA NPs were approximately 250-300 nm in diameter with or without R848. Loading of PUUC on the PLGA NP surface resulted in some aggregation, increasing the average hydrodynamic size to ~800 nm. Zeta potential of particles at pH 7.4 was measured to be ~+30 mV for MPs and NPs without PUUC. Loading with PUUC reduced NP zeta potential to ~+22 mV. For both MPs and NPs, 100% loading of PUUC adjuvant was achieved at a loading level of 10 μg/mg particles.

**Figure 1.**
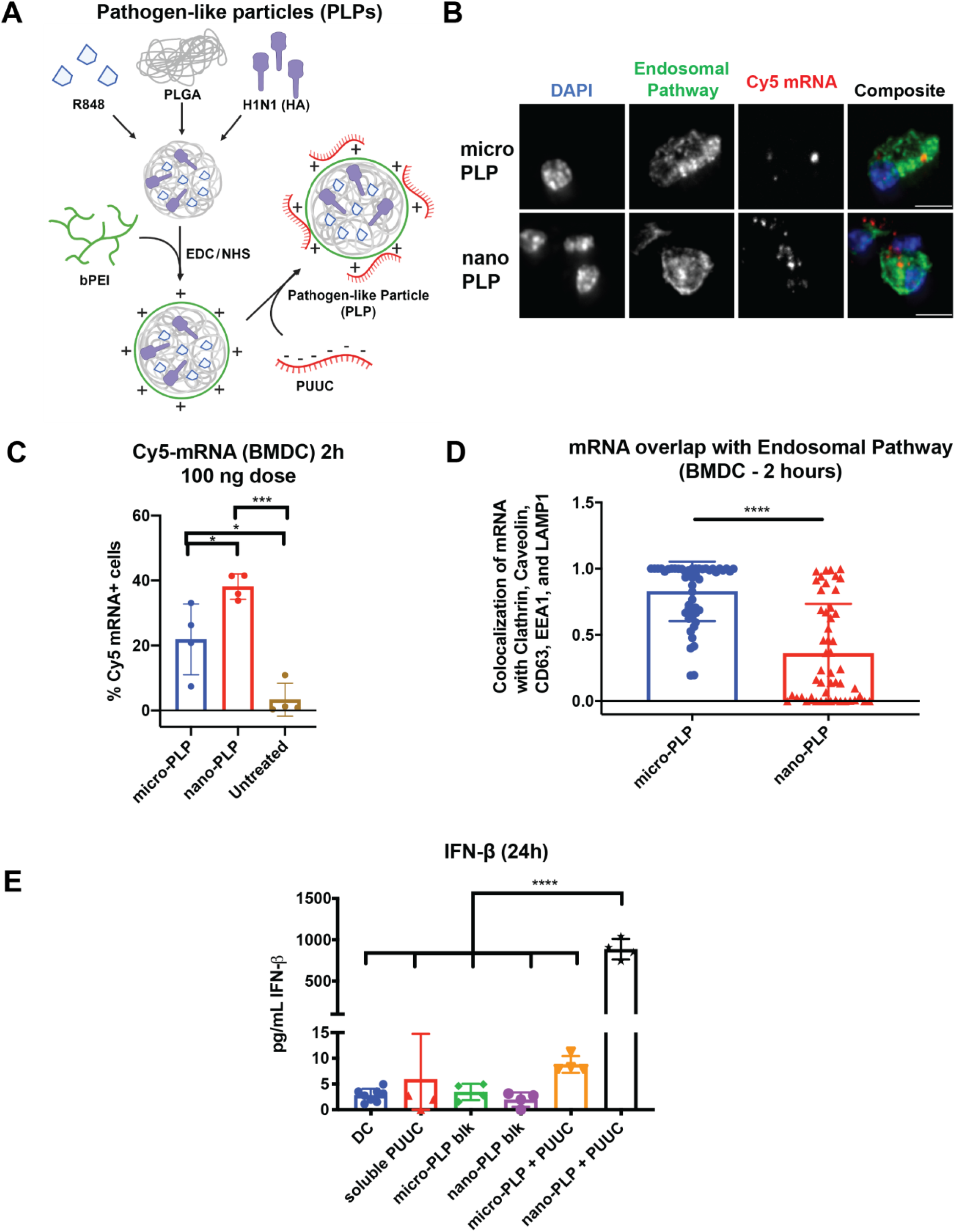
Nano-PLPs enable delivery of functional RIG-I adjuvant (PUUC). **A)** Schematic of micro-PLP or nano-PLP design. **B)** mBMDC were treated with PLGA-PEI microparticles or nanoparticles with fluorescent luciferase mRNA (red) and stained for intracellular compartments (clathrin, caveolin, CD63, EEA1, LAMP1: green). **C)** Uptake of fluorescent luciferase mRNA delivered by micro-PLPs or nano-PLPs (particle dose = 133 μg/mL), **D)** Endosomal escape was quantified by calculating Manders’ M1 overlap coefficient of mRNA with endosomal compartments (n=50 cells/treatment, scale bar = 10μm). **E)** mBMDC activation (300,000 cells/well) with soluble PUUC (500 ng/mL) or with PUUC in soluble form, micro-PLPs, or nano PLPs (150 ng/mL,17 ug/mL particle dose) after 24 hours (n=4). Error bars represent SD of the mean. For comparisons between two groups, statistical differences were determined by the Mann-Whitney test. For comparisons between more than two groups, statistical differences were determined by one-way ANOVA followed by Tukey’s test for multiple comparisons *P< 0.05,***P<0.001 ****P <0.0001.

**Table 1.** Characterization of nanoparticle formulations with and without flu antigen (HA), R848 (R848) and RIG-I adjuvant (PUUC).

| Formulation | Diameter (nm) | Zeta Potential (mV) | HA Loading (µg/mg) | PUUC Loading (µg/mg) (Efficiency %) | R848 Loading (µg/mg) (Efficiency %) |
| --- | --- | --- | --- | --- | --- |
| NP-Blank | 290.2 ± 30.27 | 30.7 ± 0.6 | - | - | - |
| MP-Blank | 1431 ± 271.7 | 32.5 ± 1.6 | - | - | - |
| NP-PUUC | 737.3 ± 121.7 | 21 ± 5.43 | - | 10.0 (100%) | - |
| MP-PUUC | 5237 ± 30.66<sup>†</sup> | -17.8 ± 7.02 | - | 10.0 (100%) | - |
| NP-R848 | 269.2 ± 31.10 | 29.1 ± 2.3 | - | - | 6.48 (64.8%) |
| NP-HA | 276.8 ± 4.74 | 30.4 ± 1.3 | 1.00 | - | - |
| NP-HA+R848 | 290.5 ± 12.43 | 33.4 ± 0.7 | 1.00 | - | 1.06 (60.8%) |
| NP-R848-PUUC | 710.5 ± 82.3 | 23.9 ± 8.07 | - | 10.0 (100%) | 6.48 (64.8%) |
| NP-R848-HA-PUUC | 768.0 ± 87.7 | 20.5 ± 4.34 | 1.00 | 10.0 (100%) | 1.06 (60.8%) |

<sup>†</sup>DLS measurements for MP-PUUC yielded a very high polydisperse index of 1.000, containing high amounts of large particles, indicating that the addition of PUUC likely aggregated the MPs. Analysis of the size of MP-PUUC by DLS may not be appropriate due to this condition.

We proceeded to assess uptake and endosomal escape of micro-PLPs and nano-PLPs. In order to evaluate cytoplasmic delivery efficacy, we stained for markers of various intracellular endo/lysosomal compartments, which included clathrin, caveolin, CD63, early endosomal marker 1 (EEA1), and lysosome-associated membrane protein 1 (LAMP1). Images revealed a higher amount of fluorescent RNA in nano-PLPs in comparison to micro-PLPs (n=50). In addition, PLPs were not all co-localized with endo/lysosomal compartments (**Figure 1B**). The median fluorescence intensity of mBMDCs treated with PLPs with fluorescent mRNA showed equivalent uptake by micro-PLP and nano-PLP formulations (**SI Figure 1B**). A significantly higher percentage of cells, however, contained RNA after nano-PLP treatment when compared to micro-PLP treatment after 2 hours (**Figure 1C**). To quantify endosomal escape, we measured the Manders’ overlap coefficient of RNA to endo/lysosomal compartments for micro-PLPs and nano-PLPs. On average, RNA delivered with nano-PLPs had lower colocalization to intracellular compartments. This suggested that nano-PLPs are more efficient at escaping into the cytosol (**Figure 1D**), compared to micro-PLPs. We then compared interferon-β secretion from mBMDCs after treatment with soluble PUUC, PLPs without adjuvant, and nano and micro-PLPs with PUUC loaded on the particle surface. Soluble adjuvant and microparticles or nanoparticles without adjuvant induced no IFN-β secretion. Microparticles with PUUC induced very low IFN-β secretion in wild-type mBMDCs and induced no IFN-β secretion in RIG-I^−/−^ mBMDCs (**SI Figure 1C**). Meanwhile, the nanoparticle with PUUC induced considerable production and secretion of interferon-β (**Figure 1E**). Nanoparticles with PUUC substituted for the double-stranded RNA analog poly(I:C) did not induce significant IFN-β secretion from wild-type mBMDCs, which indicates that the strong interferon response is specific for the combination of the delivery system and the adjuvant (**SI Figure 1D**). These studies motivated the exclusive use of the PLGA-PEI nanoparticle (from here on referred to only as NP) for all subsequent immune response studies.

### Nano-PLP delivered RIG-I agonist (PUUC) broadens the innate immune response induced by the TLR7 agonist R848 in mouse BMDCs

We next evaluated if combination of PUUC and R848 adjuvants induce stronger innate immune responses in GM-CSF differentiated mouse BMDCs as compared to adjuvant alone. In all experiments, PUUC and R848 adjuvants were co-delivered in a single nanoparticle formulation. Nanoparticles with R848 adjuvant induced TNF-α production, which was neither enhanced nor ablated with the co-delivery of PUUC adjuvant. A strong IL-1β response was induced by treatment with nanoparticles with R848 from mouse BMDCs. Co-delivery of R848 and PUUC adjuvants on nanoparticles resulted in a 50% decrease in IL-1β production from mouse BMDCs when compared to R848 alone. (**Figure 2B**). Nanoparticles with R848 induce very low levels of IFN-β when delivered alone, while the nanoparticles with PUUC induce high production of IFN-β from mouse BMDCs as shown previously. When R848 and PUUC are delivered in combination on nanoparticles, high IFN-β production is maintained (**Figure 2C**). We then measured if CXCL10, a protein whose synthesis is dependent on interferon production, was also upregulated downstream. Nanoparticles with PUUC induce high levels of CXCL10 production, while nanoparticles with R848 induce half the amount of CXCL10 production as PUUC. When PUUC and R848 are delivered in combination on nanoparticles, CXCL10 production in mouse BMDCs is comparable to when PUUC is delivered alone.

**Figure 2.**
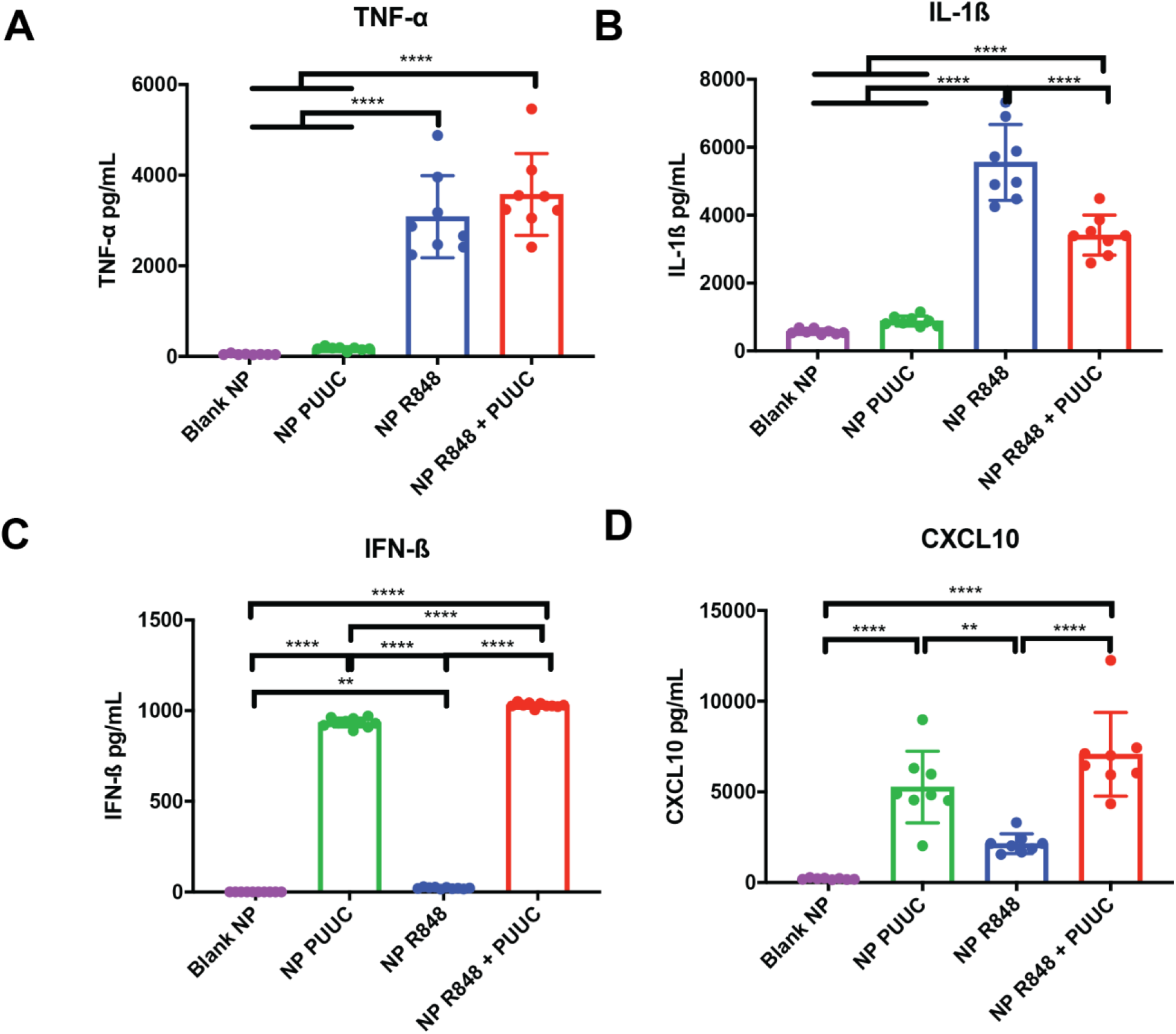
Nanoparticle co-delivery of PUUC with R848 broadens the innate immune response in GM-CSF differentiated murine BMDCs. BALB/c murine BMDCs cultured with 20 ng/mL GM-CSF were treated with blank nanoparticles (12.3 µg /mL) without adjuvant, nanoparticles with R848 (80 ng/mL), nanoparticles with PUUC (250 ng/mL), or nanoparticles co-loaded with both R848 and PUUC. Comparison of **A)** TNF-α, **B)** IL-1β, **C)** IFN-β, and **D)** CXCL10 secretion from murine BMDCs after R848 treatment with or without PUUC (n=8). Error bars represent SD of the mean. Statistical significance was determined by one-way ANOVA followed by Tukey’s test for multiple comparisons. **P≤0.01,****P≤0.0001.

### R848 + PUUC induces an additive immune response in human PBMCs and a synergistic immune response in plasmacytoid DCs (pDCs)

To investigate if the R848-PUUC adjuvant combination has relevance in humans, we evaluated innate immune activation in human PBMCs and the CAL-1 pDC line, which is derived from blastic natural killer cell lymphoma and is phenotypically similar to human plasmacytoid dendritic cells.^32^ Both NP-R848 and NP-PUUC induced IFN-α and IFN-β responses in human PBMCs, while NP-R848-PUUC induced an additive ΙFN-α and IFN-β response (**Figure 3A-B**). TNF-α responses were not significantly higher than control for the NPs with R848 or PUUC alone, but NP-R848-PUUC induced a higher TNF-α response in human PBMCs (**Figure 3C**). We proceeded to evaluate the ΙFN-α and IFN-β response in both mouse and human pDCs. A mBMDC culture differentiated with human FLT-3 was used to obtain cells with a pDC phenotype (B220+ PDCA1+). While NP-R848 and NP-PUUC induced low levels of IFN-β production, NP-R848-PUUC induced IFN-β levels that were synergistically higher than each of the individual adjuvants (**Figure 4A**). We found that PUUC delivery did not induce IFN-α or IFN-β in CAL-1 pDCs when delivered in the absence of R848 (**Figure 4B-C**). Co-delivery of PUUC with R848, however, induced a synergistic IFN-α in a dose-dependent manner. R848 induced IFN-β production, which was doubled with co-delivery of a dose of 200 ng/mL PUUC and quadrupled with co-delivery of a dose at 500 ng/mL PUUC. The R848 treatment induced TNF-α production from CAL-1 pDCs, which was not enhanced by concurrent treatment with PUUC (**SI Figure 2**).

**Figure 3.**
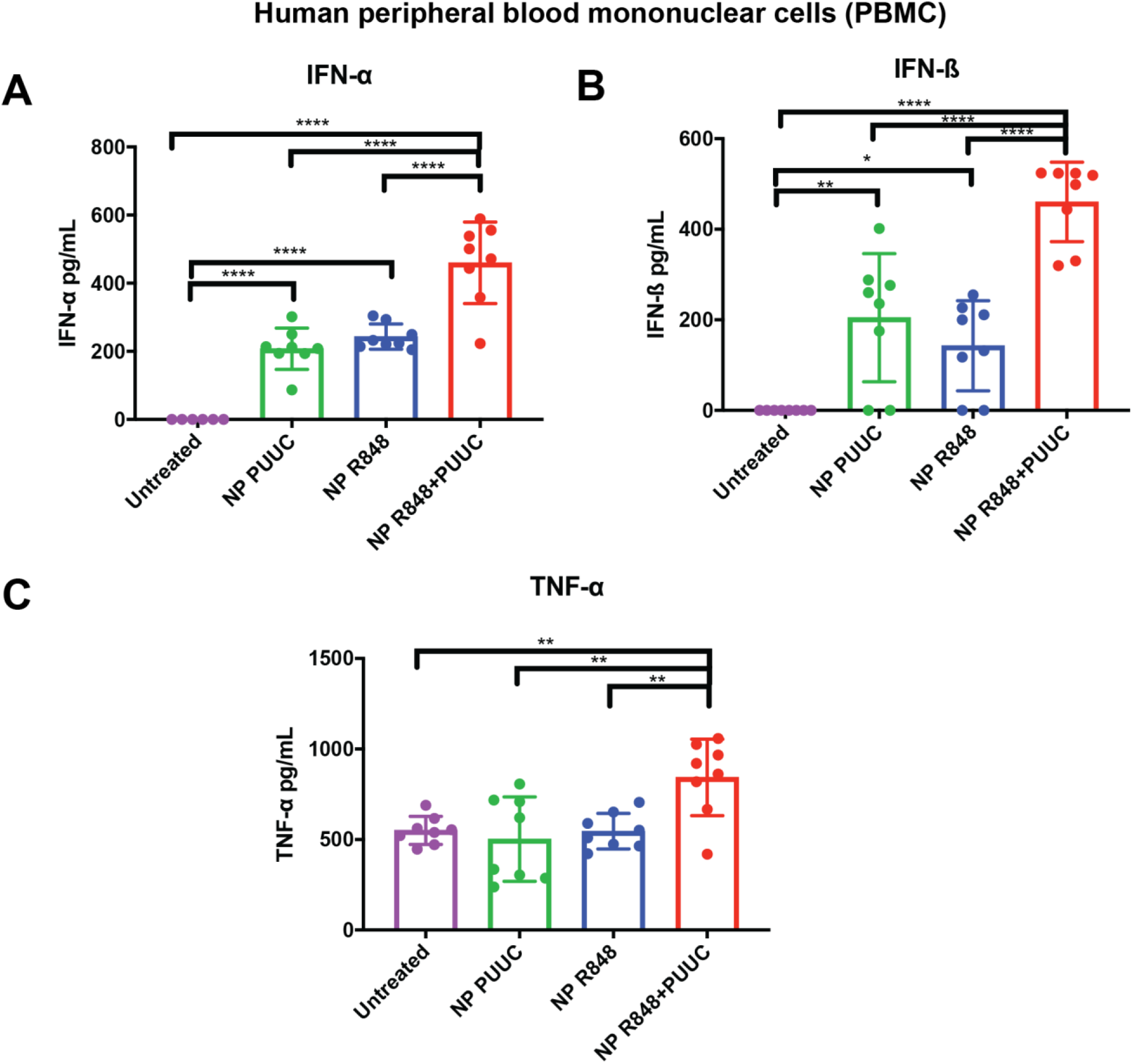
Co-delivery of R848 and PUUC in nanoparticles results in an additive innate immune response from human PBMCs. PBMCs were treated with nanoparticles (12.3 µg/mL) loaded with R848 adjuvant (80 ng/mL) and PUUC adjuvant (500 ng/mL). **A)** IFN-α, **B)** IFN-β, **C)** TNF-α were measured 24 hours after PBMC activation. In all experiments, dual delivery was performed with a single nanoparticle system. Outliers were identified by the ROUT method and removed. Statistical significance was evaluated with one-way ANOVA followed by Tukey’s test for multiple comparisons. *P≤0.05, **P≤0.01, ****P≤0.0001.

**Figure 4.**
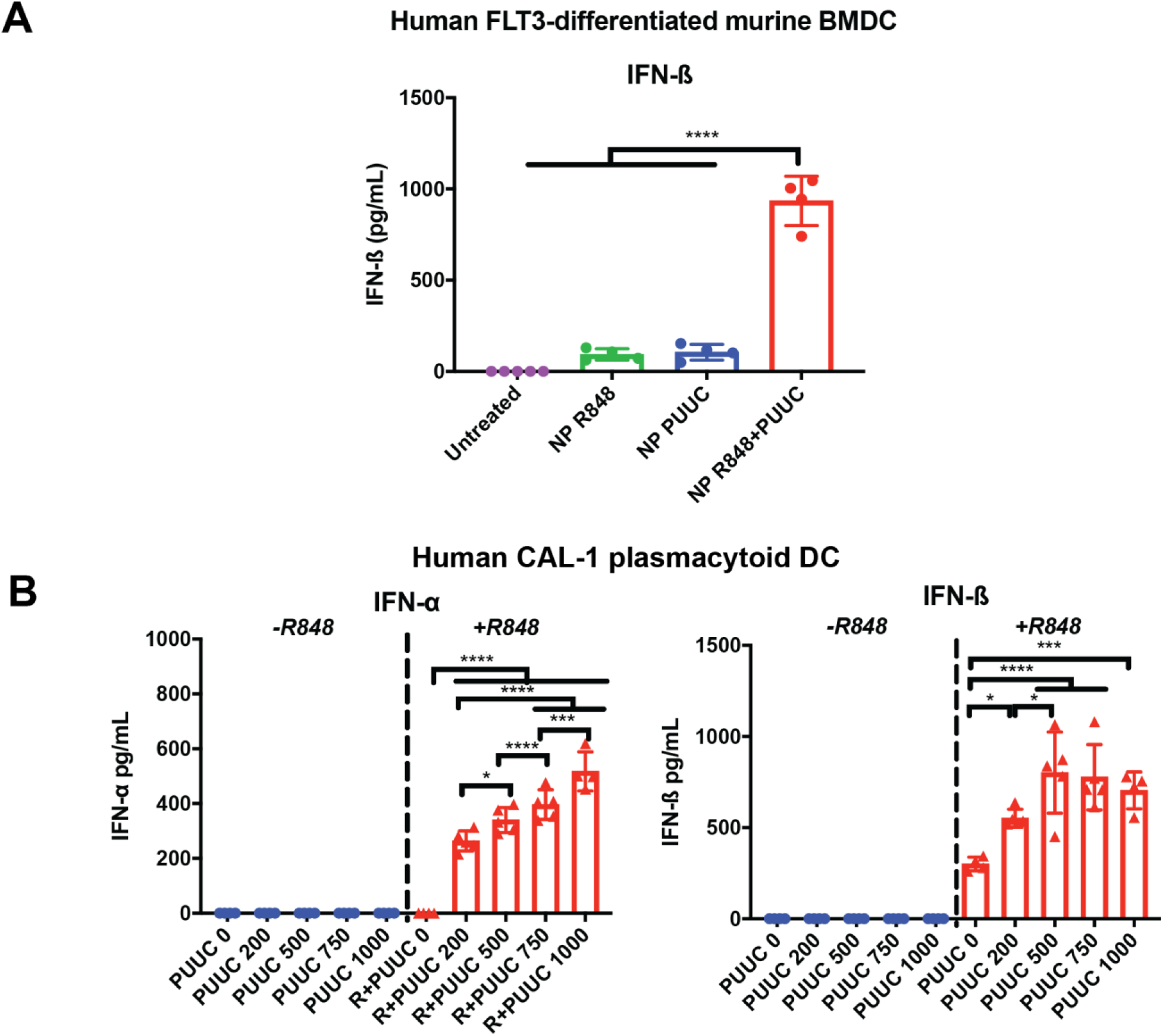
Co-delivery of R848 and PUUC in nanoparticles results in a synergistic interferon response from murine and human plasmacytoid DCs. **A)** Human FLT3-differentiated murine BMDCs were treated with nanoparticles (12.3 µg/mL) loaded with R848 adjuvant (80 ng/mL) and PUUC adjuvant (500 ng/mL) and **B-C)** CAL-1 human pDCs were treated with nanoparticles (86.5 µg/mL) loaded with R848 adjuvant (561 ng/mL) and PUUC adjuvant of doses ranging from 0-1000 ng/mL. IFN-α or IFN-β levels were measured 24 hours after pDC activation. In all experiments, dual delivery was performed with a single nanoparticle system. Nanoparticle mass was fixed across all PUUC doses. Statistical significance was evaluated with one-way ANOVA followed by Tukey’s test for multiple comparisons. *P<0.05,*** P≤0.001, ****P≤0.0001. For IFN-α, all - R848 groups are statistically different from +R848 with non-zero doses of PUUC (P≤0.0001). For IFN-β, all - R848 groups are significantly different from all +R848 groups (P≤0.0001).

### Combinatorial delivery of PUUC, R848, and a flu-HA antigen enhances antigen-specific T cell responses *in vivo*

We proceeded to evaluate if enhancements in the innate immune response would translate to improved adaptive immune responses to flu vaccines. Mice were vaccinated with nanoparticles that encapsulate H1N1 flu antigen, R848, and PUUC. The mice were given a booster dose 28 days after the initial vaccination and tissues were analyzed one week after boost (**Figure 5A**). Nanoparticles with H1N1 antigen, R848, and PUUC upregulated CD8+ T cell populations in the lung significantly more than nanoparticles with H1N1 antigen and either R848 or PUUC (**Figure 5B**). Antigen-specific CD4 T-cell populations in the spleen were increased in mice vaccinated with H1N1 antigen, and PUUC (**Figure 5C**, **SI Figure 3**). In the popliteal lymph nodes, nanoparticles with H1N1 antigen, R848, and PUUC boosted the percentage of high B220+ expressing cells more than any other vaccine regimen (**SI Figure 4A**). Mice vaccinated with H1N1 antigen alone, however, produced significantly more H1N1-specific IgG1 antibodies than mice vaccinated with H1N1 antigen and single or dual adjuvants. There were no significant differences in H1N1-specific IgG2a titers (**SI Figure 4B**). Memory cell populations in the spleen were unaffected by the delivery of adjuvants. (**SI Figure 4C-E**). Splenocytes were harvested and then restimulated with 0.5 μg H1N1 antigen (2.5 μg/mL) for 6 hours. After restimulation, CD3+CD4+ cells from mice vaccinated with H1N1 antigen, R848, and PUUC produced significantly more IFN-γ, IL-4, and TNF-α than splenocytes from mice vaccinated with either antigen alone or antigen with a single adjuvant (**Figure 5D**). The H1N1 antigen/R848/PUUC combination also enhanced the percentage of CD3+ CD8+ T-cells expressing IFN-γ, IL-4, and TNF-α after restimulation (**SI Figure 4F**). A significantly higher percentage of CD3+ CD4+ and CD3+ CD8+ T-cells treated with H1N1 antigen, R848, and PUUC produced both IFN-γ and TNF-α when compared to all other groups (**SI Figure 4G**). Interestingly, similar to that seen in viral infections, including SARS-CoV-2,^33,34^ which carries TLR7 and PUUC ligands together, there were an overall lower number of T-cells in the spleen of the dual-adjuvant treated animals, indicating some virus-like lymphopenia. Similarly in the restimulated splenocytes, the relative amount of CD4+ and CD8+ T-cells from mice vaccinated with dual adjuvants was significantly lower (indicating potential overstimulation of the T cells) when compared to all other treatment groups (**SI Figure 5A-B**).

**Figure 5.**
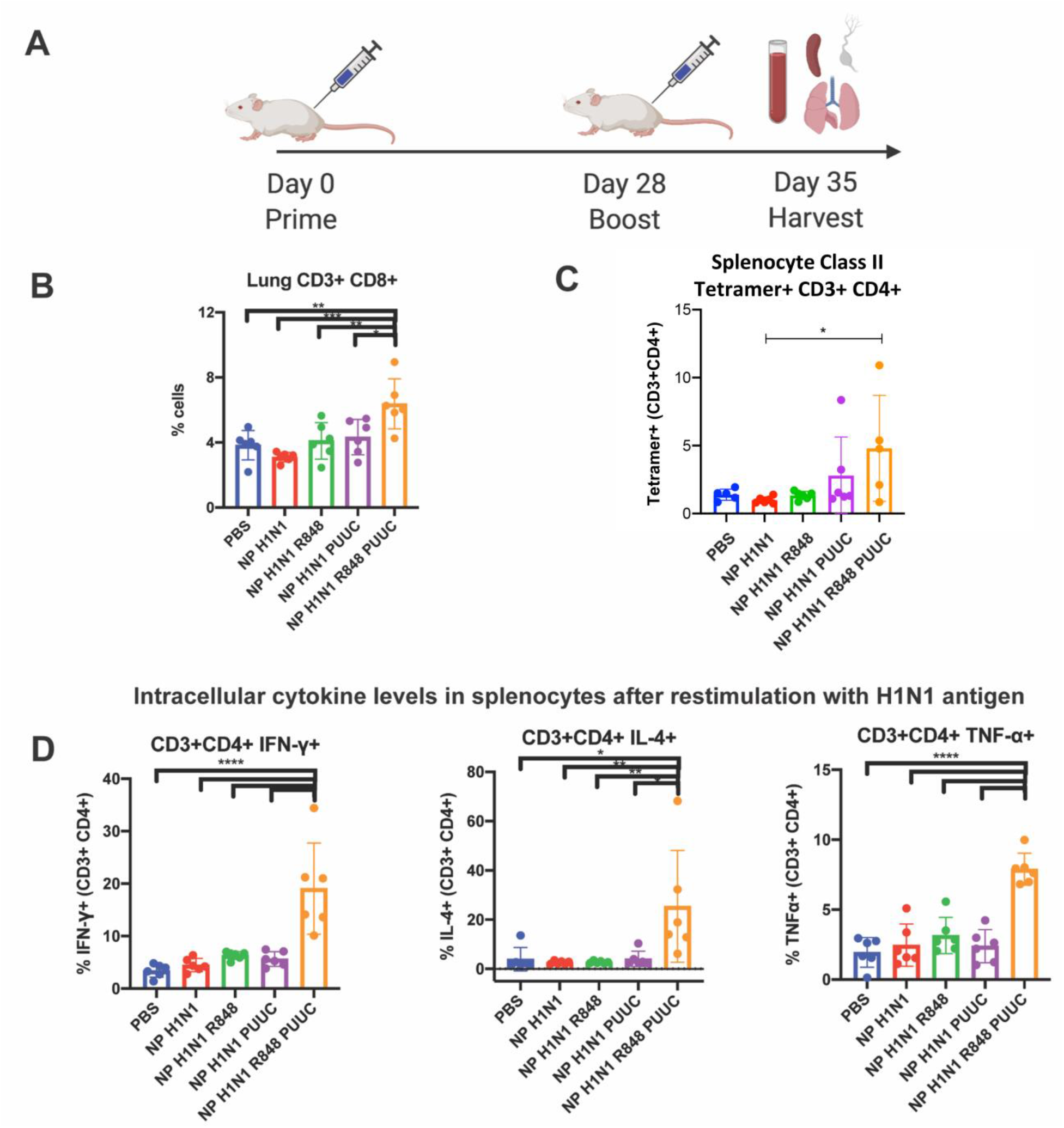
TLR7 adjuvant (R848) and RIG-I adjuvant (PUUC) induce enhanced T-cell proliferation and recall responses to flu antigen in nanoparticles. **A)**BALB/c mice were vaccinated by intramuscular injection into both quadricep muscles on day 0 and day 28. Treatment groups included PLGA nanoparticles with H1N1 antigen (NP Η1Ν1), H1N1 antigen and R848 (NP H1Ν1 R848), H1N1 antigen and PUUC (NP Η1Ν1 PUUC), and H1N1 antigen and R848 and PUUC (NP H1Ν1 R848 PUUC). Dosing per injection was as follows: PLGA NP = 0.5 mg, H1N1 = 0.5 μg, R848 = 0.525 μg, PUUC = 10 μg. Lungs, blood, spleen, and lymph nodes were harvested after 35 days. **B)** Percentage of CD3+ CD8+ cells in lung. **C)** Percentage of Class II AELLVLLEN tetramer+ CD3+ CD8-(presumably CD4+) cells in splenocytes. Intracellular levels of IFN-γ, IL-4, and TNF-α were measured in **D)** CD3+ CD4+ live splenocytes restimulated with H1N1 antigen for 6 hours. In all experiments, antigens and single or dual adjuvants were co-delivered on a single nanoparticle system. Error bars represent SD of the mean. Statistical significance was determined by one-way ANOVA followed by Tukey’s test for multiple comparisons for normal datasets. *P≤ 0.05, **P≤ 0.01, ***P≤0.001, ****P≤0.0001.

## DISCUSSION

We developed a PLP that delivers functional RIG-I adjuvant (PUUC) *in vitro* and *in vivo* and demonstrated that co-delivery with the TLR7 adjuvant R848 results in broadened or synergistic immune responses. Flu viruses are difficult to vaccinate against because of their rapidly mutating epitopes and their induction of non-structural protein expression, which suppresses the host innate immune response.^35^ In subunit vaccines, low immunogenicity is a serious problem. Many individual adjuvants have been studied for enhancement of flu vaccine immunogenicity.^9,10,23,36^ A few adjuvant combinations have been studied in conjunction with flu, but combinations of TLR and RLR adjuvants have not thoroughly been investigated.^13,14^

To deliver a combination of TLR and RLR adjuvants, we synthesized PLPs that deliver both protein antigen and adjuvants (e.g., R848) within the polymer matrix and charged adjuvants on the particle surface (e.g., PUUC). Similar pathogen-like particles have successfully delivered encapsulated TLR4 and TLR7 adjuvants with charged TLR9 adjuvants, which induced *in vitro* and *in vivo* synergistic immune responses.^37^ We tested if microparticles (~1.5 μm diameter) or nanoparticles (~250 nm diameter) could induce IFN-β responses after presentation with PUUC. Only the nanoparticles with PUUC induced a robust IFN-β response in murine GM-CSF differentiated BMDCs. Although the nanoparticles with PUUC aggregated, we ruled out size as a trigger for Type I interferon production by demonstrating that the interferon response to nanoparticles loaded with poly(I:C), a double-stranded RNA analog, was absent. Consistent loading efficiency of PUUC across nanoparticles with and without antigen or adjuvant suggest insignificant differences in the charge of PUUC-loaded formulations. We hypothesized that the different responses were due to differences in cellular uptake and endosomal escape of the microparticles and nanoparticles – two processes that are essential for cytosolic delivery to RLRs. Although the average amount of mRNA delivered per cell was equal for micro-PLPs and nano-PLPs, we observed that a higher percentage of cells were transfected with nano-PLPs. Nanoparticles also escaped the endolysosomal pathway into the cytosol more efficiently than microparticles. For cationic lipid nanoparticles, size plays a role in determining a particle’s ability to bend and destabilize the plasma membrane for endosomal rupture.^38^ With cationic polymer nanoparticles, such as our PLP system, endosomal escape is hypothesized to be driven by the proton sponge effect.^39^ The monolayer of branched PEI on our PLPs, which has exposed secondary or tertiary amines, enhances the proton sponge effect. Toxicity is also lower with PLPs in comparison to free PEI. As an endosome matures into a lysosome, the compartmental pH drops and the amines of the PEI on the PLP surface become protonated. In turn, this drives osmotic flux into the lysosome and subsequently bursts the compartment. Because particle mass and adjuvant dose are held constant in experiments, there is a higher total surface area with nano-PLPs in comparison to micro-PLPs. Therefore, we speculate that the enhanced endosomal escape of nano-PLPs is due to a higher amount of PEI available to drive osmotic flux for rupture of lysosomes. Our findings justified the use of nanoparticles instead of microparticles for co-delivery of R848 and PUUC, which can also present adjuvants that activate endosomal TLRs.^31^

Co-delivery of PUUC, a RIG-I agonist, and R848, a hydrophobic TLR7/8 agonist, produces a diverse cytokine response from murine GM-CSF-differentiated BMDCs. This response is expected from stimulation of RLRs, which phosphorylate IRF3 and IRF7 to promote Type I interferon expression, and stimulation of TLR7, which triggers a significant TNF-α response through MyD88-dependent NF-kB activation.^40,41^ TLR7 stimulation by R848 stimulates production of the procytokine pro-IL-1β.^42^ While R848 stimulation alone induces low levels of mature IL-1β, it is likely that the cationic PLPs amplify activation of inflammasome complexes by disrupting membrane potential.^43^ We showed that the loading of anionic PUUC onto PLPs with R848 reduces zeta potential, which may explain why IL-1β production in PLPs with both R848 and PUUC is dampened. NF-κΒ also downregulates inflammasome activation, so enhanced NF-κΒ activation by PUUC may also be contributing to the slight dampening of the IL-1β response.^44^ In human FLT3-differentiated murine BMDCs and the human CAL-1 pDC cell line, nanoparticle co-delivery of R848 and PUUC induced synergistic cytokine outputs of IFN-α and IFN-β. The interferon response is critical for flu protection, as successful flu infection is facilitated by viral nonstructural protein-mediated inhibition of the interferon induction signaling cascade. Unlike in murine GM-CSF differentiated BMDCs, which generated an IFN-α response with PUUC alone, human CAL-1 pDCs required co-delivery of both R848 and PUUC to produce IFN-α. TLR7-mediated synergy has also been observed in CAL-1 pDCs when two unique TLR7 adjuvants have been delivered together.^45^ It is possible that CAL-1 pDCs require the activation of multiple pattern recognition receptors (PRRs) to produce IFN-α. Unlike IFN-α, IFN-β can be produced by CAL-1 pDCs either with R848 alone or with PUUC in a dose-dependent manner. Because PUUC loading density on NPs was increased to maintain the same R848 dose across groups, the loading density may also play a role in the magnitude of the interferon response. It is likely that R848 is upregulating RIG-I expression in the cytosol in pDCs, which upregulates Type I interferon production by PUUC activation.^46^ Conventional BMDCs, which express RLRs constitutively, do not exhibit this phenemonon.^22^ These findings motivate further investigation of signaling crosstalk in CAL-1 pDCs, which may identify new therapeutic targets that induce strong antiviral responses in pDCs.

Although antibody-mediated immunity is the main correlate of protection for flu, it has been suggested that cell-mediated immunity could enhance protection.^47^ CD8+ T-cells rapidly proliferate in response to antigen; effector cell populations subsequently contract, which leave a memory T-cell population that can proliferate and secrete cytokines after restimulation. Indeed, T cell cytokine and Granzyme B responses correlate with influenza resistance in vaccinated seniors, while antibody titers did not correlate to protection from flu.^48^ Furthermore, CD8+ T cells have been implicated in responses to heterotypic influenza infections, possibly by recognizing conserved internal influenza peptides.^49,50^ This cross-reactivity may explain unexpected mild illnesses in the 2009 pandemic H1N1 virus.^7,51^

While the combination of R848 and PUUC did not enhance antibody-mediated immunity, it enhanced multiple aspects of cellular immunity. In the lung, CD8+ T-cell populations increased after vaccination with H1N1 antigen, R848, and PUUC without altering the percentage of tissue-resident T-cells. Higher percentage of H1N1 tetramer positive CD4+ T-cells in the splenocytes were also observed with the vaccine and adjuvant combination. A significant increase in intracellular levels of IFN-γ, IL-4, and TNF-α - three cytokines critical for an antiviral response upon re-exposure to flu - was observed in splenocytes activated with R848 and PUUC - a phenomenon not observed after treatment with individual adjuvants. The enhanced cellular response may be attributed to the combination of TLR7-mediated upregulation of TNF-α and RIG-I-mediated upregulation of Type I interferon and interferon-stimulated genes, such as CXCL10. TNF-α increases vascular permeability and increases cellular adhesion molecule expression, which aids extravasation of leukocytes into inflamed tissues.^52^ RIG-I signaling is a critical event that triggers Type I interferon production and upregulates interferon stimulated genes, such as CXCL10, to promote migratory function in both activated T-cells and dendritic cells.^53,54^ Cross-presentation is also enhanced by both TLR7 and RIG-I activation.^55,56^ Therefore, synergistic cellular immunity through concurrent TLR7 and RIG-I activation may be driven by enhanced recruitment and priming of both effector and antigen-presenting cells. We did not observe changes in effector or memory T-cell populations, but this is reasonable when considering that samples were analyzed one week after boost. Typically, it takes weeks to months for memory T-cell populations to emerge after vaccination.^57^

Also, both before and after antigen restimulation, the dual adjuvant splenocyte group showed relatively reduced proportions of CD4 and CD8 T-cells. In splenocytes stained before antigen restimulation, this cell death is consistent with other reports of TLR7- or RIG-I-related lymphopenia,^58–60^ and is an interesting feature of a delivery system that models these virus-linked innate immune signals. Indeed, T-cell lymphopenia itself has been observed with influenza illness.^61,62^ Similarly, the presence of T-cell lymphopenia in the restimulated splenocytes suggests that T-cells in that group could be overstimulated or highly sensitive. While neither adjuvant significantly caused this effect alone, the observed lymphopenia is possibly attributable to the increased type I interferon production associated with dual TLR7 and RIG-I stimulation, as shown in our *in vitro* experiments. Again, this may be attributable to upregulation and overexpression of RIG-I in DCs by TLR7 ligation with R848, followed by RIG-I stimulation by PUUC leading to increased interferon production.^46^ Additionally, the overexpression and stimulation of the RIG-I pathway may be contributing to reduced T-cell proliferation and apoptosis, as has been associated with lymphopenia in patients with dermatomyositis.^60^ Though TLR7-associated transient lymphopenia has been studied for decades,^63^ the role of RIG-I-related lymphopenia is less clear. Given that TLR7 and RIG-I ligation induce broadened and synergistic proinflammatory cytokine secretions, and that such cytokines may be a cause of lymphopenia in respiratory diseases like COVID-19 and influenza,^64,65^ further investigation into the combined effects of this adjuvant combination on T-cell viability and PRR signaling is warranted.

In summary, we have demonstrated that a hydrophobic TLR7 adjuvant and a RIG-I adjuvant make up a potent adjuvant combination that drives broadened and synergistic immune responses. Future studies should determine the precise mechanism in which antibody-mediated responses were not strengthened with the adjuvant combination, a phenomenon also observed in a previous report using particles with co-encapsulated HA antigen and R848.^14^ In this report, the separation of antigens and adjuvants into separate nanoparticles revived the humoral response. The mechanism for this is unknown, but it is possible that antigen-processing and presentation to Tfh cells by lymph node B cells is hampered by co-loaded adjuvants. In addition, we propose future study of T-cell memory at a later timepoint after vaccination and evaluation if strong cell-mediated responses lead to enhanced vaccine efficacy in challenge studies. Because of their relevance to influenza infections, the response of antigen-specific T cell populations in the lung should also be explored in relation to combinatorial TLR7 and RIG-I ligation. Our results motivate the development of immune adjuvant systems that engage both TLRs and cytosolic innate immune receptors. An intriguing future vaccine design could be a PLP system that combines TLR7 and RIG-I adjuvants with nucleoprotein, an antigen that is associated with strong cellular immune responses.^66^ Targeted delivery to subsets of innate immune cells that synergistically respond to adjuvant combinations may pave the way to more robust vaccines and antiviral therapies.

## Supporting information

Supplemental Information - Version 2

## ACKNOWLEDGEMENTS

We thank Joscelyn Mejias for advice with image analysis, Alexander Beach for the culturing of human FLT3-differentiated murine BMDCs, Melissa Kemp for allowing generous use of her lab’s BioPlex 200 Luminex instrument, James Bowen and Kendra Quicke for their assistance with early studies with the PUUC adjuvant, Takahiro Maeda and Dennis Klinman for their gift of CAL-1 pDCs, Michael Gale Jr. for the gift of RIG-I^−/−^ mice, and Philip Santangelo for provision of mRNA and advice on the endosomal escape assay. We acknowledge the use of the Engineered Biosystems Building Physiological Research Lab for animal experiments, Optical Microscopy Core for microscopy experiments, Cellular Analysis Core for flow cytometry experiments, and Biopolymer Characterization Core for the preparation of nanoparticles. The graphical abstract, particle assembly schematic, and vaccine timeline images were created with BioRender.com. We acknowledge funding from the Georgia Tech Foundation, the Georgia Clinical and Translational Science Alliance for support of collaborative efforts between Mehul Suthar and Krishnendu Roy, NIH grant U01-AI124270-02, the Georgia Tech President’s Undergraduate Research Award for support of M. Cole Keenum, Karl Dasher and family for their support of M. Cole Keenum through the Georgia Tech Petit Scholars program, the Cell Manufacturing Technologies (CMaT) Research Experiences for Undergraduates program for support to Chinwendu Chukwu, and the Robert A. Milton Chaired Professorship to Krishnendu Roy.

## REFERENCES

1 Ed., K. Influenza Pandemics of the 20th Century. Emerg Infect Dis 12(1), 9–14 (2006).

2 Johnson, N. P. A. S. a. J. M. Updating the Accounts: Global Mortality of the 1918-1920 “Spanish Influenza Pandemic”. Bulletin of the History of Medicine 76, 105–115, doi:doi:10.1353/bhm.2002.0022 (2002).

3 Paget, J. et al. Global mortality associated with seasonal influenza epidemics: New burden estimates and predictors from the GLaMOR Project. J Glob Health 9, 020421, doi:10.7189/jogh.09.020421 (2019).

4 Lansbury, L. E. et al. Effectiveness of 2009 pandemic influenza A(H1N1) vaccines: A systematic review and meta-analysis. Vaccine 35, 1996–2006, doi:10.1016/j.vaccine.2017.02.059 (2017).

5 Heaton, N. S., Sachs, D., Chen, C. J., Hai, R. & Palese, P. Genome-wide mutagenesis of influenza virus reveals unique plasticity of the hemagglutinin and NS1 proteins. Proc Natl Acad Sci U S A 110, 20248–20253, doi:10.1073/pnas.1320524110 (2013).

6 Clemens, E. B., van de Sandt, C., Wong, S. S., Wakim, L. M. & Valkenburg, S. A. Harnessing the Power of T Cells: The Promising Hope for a Universal Influenza Vaccine. Vaccines (Basel) 6, doi:10.3390/vaccines6020018 (2018).

7 Pizzolla, A. & Wakim, L. M. Memory T Cell Dynamics in the Lung during Influenza Virus Infection. J Immunol 202, 374–381, doi:10.4049/jimmunol.1800979 (2019).

8 Ho, N. I., Huis In 't Veld, L. G. M., Raaijmakers, T. K. & Adema, G. J. Adjuvants Enhancing Cross-Presentation by Dendritic Cells: The Key to More Effective Vaccines? Front Immunol 9, 2874, doi:10.3389/fimmu.2018.02874 (2018).

9 van der Most, R. G. et al. Long-Term Persistence of Cell-Mediated and Humoral Responses to A(H1N1)pdm09 Influenza Virus Vaccines and the Role of the AS03 Adjuvant System in Adults during Two Randomized Controlled Trials. Clin Vaccine Immunol 24, doi:10.1128/CVI.00553-16 (2017).

10 Vesikari, T., Forsten, A., Arora, A., Tsai, T. & Clemens, R. Influenza vaccination in children primed with MF59-adjuvanted or non-adjuvanted seasonal influenza vaccine. Hum Vaccin Immunother 11, 2102–2112, doi:10.1080/21645515.2015.1044167 (2015).

11 Goldinger, S. M. et al. Nano-particle vaccination combined with TLR-7 and −9 ligands triggers memory and effector CD8(+) T-cell responses in melanoma patients. Eur J Immunol 42, 3049–3061, doi:10.1002/eji.201142361 (2012).

12 Moody, M. A. et al. Toll-like receptor 7/8 (TLR7/8) and TLR9 agonists cooperate to enhance HIV-1 envelope antibody responses in rhesus macaques. J Virol 88, 3329–3339, doi:10.1128/JVI.03309-13 (2014).

13 Goff, P. H. et al. Synthetic Toll-like receptor 4 (TLR4) and TLR7 ligands as influenza virus vaccine adjuvants induce rapid, sustained, and broadly protective responses. J Virol 89, 3221–3235, doi:10.1128/JVI.03337-14 (2015).

14 Kasturi, S. P. et al. Programming the magnitude and persistence of antibody responses with innate immunity. Nature 470, 543–547, doi:10.1038/nature09737 (2011).

15 Hammerbeck, D. M. et al. Administration of a dual toll-like receptor 7 and toll-like receptor 8 agonist protects against influenza in rats. Antiviral Res 73, 1–11, doi:10.1016/j.antiviral.2006.07.011 (2007).

16 Errett, J. S., Suthar, M. S., McMillan, A., Diamond, M. S. & Gale, M., Jr. The essential, nonredundant roles of RIG-I and MDA5 in detecting and controlling West Nile virus infection. J Virol 87, 11416–11425, doi:10.1128/JVI.01488-13 (2013).

17 Bowen, J. R. et al. Zika Virus Antagonizes Type I Interferon Responses during Infection of Human Dendritic Cells. PLoS Pathog 13, e1006164, doi:10.1371/journal.ppat.1006164 (2017).

18 Weber-Gerlach, M. & Weber, F. Standing on three legs: antiviral activities of RIG-I against influenza viruses. Curr Opin Immunol 42, 71–75, doi:10.1016/j.coi.2016.05.016 (2016).

19 Zhao, L. et al. Identification of cellular microRNA-136 as a dual regulator of RIG-I-mediated innate immunity that antagonizes H5N1 IAV replication in A549 cells. Sci Rep 5, 14991, doi:10.1038/srep14991 (2015).

20 Wu, W. et al. RIG-I and TLR3 are both required for maximum interferon induction by influenza virus in human lung alveolar epithelial cells. Virology 482, 181–188, doi:10.1016/j.virol.2015.03.048 (2015).

21 Probst, P. et al. A small-molecule IRF3 agonist functions as an influenza vaccine adjuvant by modulating the antiviral immune response. Vaccine 35, 1964–1971, doi:10.1016/j.vaccine.2017.01.053 (2017).

22 Fekete, T. et al. Human Plasmacytoid and Monocyte-Derived Dendritic Cells Display Distinct Metabolic Profile Upon RIG-I Activation. Front Immunol 9, 3070, doi:10.3389/fimmu.2018.03070 (2018).

23 Kulkarni, R. R. et al. Activation of the RIG-I pathway during influenza vaccination enhances the germinal center reaction, promotes T follicular helper cell induction, and provides a dose-sparing effect and protective immunity. J Virol 88, 13990–14001, doi:10.1128/JVI.02273-14 (2014).

24 Beljanski, V. et al. Enhanced Influenza Virus-Like Particle Vaccination with a Structurally Optimized RIG-I Agonist as Adjuvant. J Virol 89, 10612–10624, doi:10.1128/JVI.01526-15 (2015).

25 Lund, J. M. et al. Recognition of single-stranded RNA viruses by Toll-like receptor 7. Proc Natl Acad Sci U S A 101, 5598–5603, doi:10.1073/pnas.0400937101 (2004).

26 Pichlmair, A. et al. RIG-I-mediated antiviral responses to single-stranded RNA bearing 5'-phosphates. Science 314, 997–1001, doi:10.1126/science.1132998 (2006).

27 Spitaels, J., Roose, K. & Saelens, X. In fluenza and Memory T Cells: How to Awake the Force. Vaccines (Basel) 4, doi:10.3390/vaccines4040033 (2016).

28 Stone, A. E. L., Green, R., Wilkins, C., Hemann, E. A. & Gale, M. RIG-I-like receptors direct inflammatory macrophage polarization against West Nile virus infection. Nature Communications 10, 3649, doi:10.1038/s41467-019-11250-5 (2019).

29 Saito, T., Owen, D. M., Jiang, F., Marcotrigiano, J. & Gale, M., Jr. Innate immunity induced by composition-dependent RIG-I recognition of hepatitis C virus RNA. Nature 454, 523–527, doi:10.1038/nature07106 (2008).

30 Santangelo, P. J. et al. Single molecule-sensitive probes for imaging RNA in live cells. Nat Methods 6, 347–349, doi:10.1038/nmeth.1316 (2009).

31 Leleux, J. A., Pradhan, P. & Roy, K. Biophysical Attributes of CpG Presentation Control TLR9 Signaling to Differentially Polarize Systemic Immune Responses. Cell Rep 18, 700–710, doi:10.1016/j.celrep.2016.12.073 (2017).

32 Carmona-Saez, P. et al. Metagene projection characterizes GEN2.2 and CAL-1 as relevant human plasmacytoid dendritic cell models. Bioinformatics 33, 3691–3695, doi:10.1093/bioinformatics/btx502 (2017).

33 Huang, C. et al. Clinical features of patients infected with 2019 novel coronavirus in Wuhan, China. The lancet 395, 497–506 (2020).

34 Yang, X. et al. Clinical course and outcomes of critically ill patients with SARS-CoV-2 pneumonia in Wuhan, China: a single-centered, retrospective, observational study. The Lancet Respiratory Medicine (2020).

35 Pichlmair, A. & Reis e Sousa, C. Innate recognition of viruses. Immunity 27, 370–383, doi:10.1016/j.immuni.2007.08.012 (2007).

36 Junkins, R. D. et al. A robust microparticle platform for a STING-targeted adjuvant that enhances both humoral and cellular immunity during vaccination. J Control Release 270, 1–13, doi:10.1016/j.jconrel.2017.11.030 (2018).

37 Madan-Lala, R., Pradhan, P. & Roy, K. Combinatorial Delivery of Dual and Triple TLR Agonists via Polymeric Pathogen-like Particles Synergistically Enhances Innate and Adaptive Immune Responses. Sci Rep 7, 2530, doi:10.1038/s41598-017-02804-y (2017).

38 Peetla, C., Jin, S., Weimer, J., Elegbede, A. & Labhasetwar, V. Biomechanics and thermodynamics of nanoparticle interactions with plasma and endosomal membrane lipids in cellular uptake and endosomal escape. Langmuir 30, 7522–7532, doi:10.1021/la5015219 (2014).

39 Akinc, A., Thomas, M., Klibanov, A. M. & Langer, R. Exploring polyethylenimine-mediated DNA transfection and the proton sponge hypothesis. J Gene Med 7, 657–663, doi:10.1002/jgm.696 (2005).

40 Yong, H. Y. & Luo, D. RIG-I-Like Receptors as Novel Targets for Pan-Antivirals and Vaccine Adjuvants Against Emerging and Re-Emerging Viral Infections. Front Immunol 9, 1379, doi:10.3389/fimmu.2018.01379 (2018).

41 Dowling, D. J. Recent Advances in the Discovery and Delivery of TLR7/8 Agonists as Vaccine Adjuvants. Immunohorizons 2, 185–197, doi:10.4049/immunohorizons.1700063 (2018).

42 Fernandez, M. V. et al. Ion efflux and influenza infection trigger NLRP3 inflammasome signaling in human dendritic cells. J Leukoc Biol 99, 723–734, doi:10.1189/jlb.3A0614-313RRR (2016).

43 Warren, E. A. & Payne, C. K. Cellular binding of nanoparticles disrupts the membrane potential. RSC Adv 5, 13660–13666, doi:10.1039/C4RA15727C (2015).

44 Liu, T., Zhang, L., Joo, D. & Sun, S. C. NF-kappaB signaling in inflammation. Signal Transduct Target Ther 2, doi:10.1038/sigtrans.2017.23 (2017).

45 Hilbert, T. et al. Synergistic Stimulation with Different TLR7 Ligands Modulates Gene Expression Patterns in the Human Plasmacytoid Dendritic Cell Line CAL-1. Mediators Inflamm 2015, 948540, doi:10.1155/2015/948540 (2015).

46 Szabo, A. et al. TLR ligands upregulate RIG-I expression in human plasmacytoid dendritic cells in a type I IFN-independent manner. Immunol Cell Biol 92, 671–678, doi:10.1038/icb.2014.38 (2014).

47 La Gruta, N. L. & Turner, S. J. T cell mediated immunity to influenza: mechanisms of viral control. Trends Immunol 35, 396–402, doi:10.1016/j.it.2014.06.004 (2014).

48 McElhaney, J. E. et al. T cell responses are better correlates of vaccine protection in the elderly. J Immunol 176, 6333–6339, doi:10.4049/jimmunol.176.10.6333 (2006).

49 Auladell, M. et al. Recalling the Future: Immunological Memory Toward Unpredictable Influenza Viruses. Front Immunol 10, 1400, doi:10.3389/fimmu.2019.01400 (2019).

50 Gotch, F., Rothbard, J., Howland, K., Townsend, A. & McMichael, A. Cytotoxic T lymphocytes recognize a fragment of influenza virus matrix protein in association with HLA-A2. Nature 326, 881–882, doi:10.1038/326881a0 (1987).

51 Tu, W. et al. Cytotoxic T lymphocytes established by seasonal human influenza cross-react against 2009 pandemic H1N1 influenza virus. J Virol 84, 6527–6535, doi:10.1128/JVI.00519-10 (2010).

52 Teijeira, A. et al. T Cell Migration from Inflamed Skin to Draining Lymph Nodes Requires Intralymphatic Crawling Supported by ICAM-1/LFA-1 Interactions. Cell Rep 18, 857–865, doi:10.1016/j.celrep.2016.12.078 (2017).

53 Liu, M. et al. CXCL10/IP-10 in infectious diseases pathogenesis and potential therapeutic implications. Cytokine Growth Factor Rev 22, 121–130, doi:10.1016/j.cytogfr.2011.06.001 (2011).

54 Dufour, J. H. et al. IFN-gamma-inducible protein 10 (IP-10; CXCL10)-deficient mice reveal a role for IP-10 in effector T cell generation and trafficking. J Immunol 168, 3195–3204, doi:10.4049/jimmunol.168.7.3195 (2002).

55 Crespo, M. I. et al. TLR7 triggering with polyuridylic acid promotes cross-presentation in CD8alpha+ conventional dendritic cells by enhancing antigen preservation and MHC class I antigen permanence on the dendritic cell surface. J Immunol 190, 948–960, doi:10.4049/jimmunol.1102725 (2013).

56 Hochheiser, K. et al. Cutting Edge: The RIG-I Ligand 3pRNA Potently Improves CTL Cross-Priming and Facilitates Antiviral Vaccination. J Immunol 196, 2439–2443, doi:10.4049/jimmunol.1501958 (2016).

57 Cui, W. & Kaech, S. M. Generation of effector CD8+ T cells and their conversion to memory T cells. Immunol Rev 236, 151–166, doi:10.1111/j.1600-065X.2010.00926.x (2010).

58 Perkins, H. et al. Therapy with TLR7 Agonists Induces Lymphopenia: Correlating Pharmacology to Mechanism in a Mouse Model. Journal of Clinical Immunology 32, 1082–1092, doi:10.1007/s10875-012-9687-y (2012).

59 Baenziger, S. et al. Triggering TLR7 in mice induces immune activation and lymphoid system disruption, resembling HIV-mediated pathology. Blood 113, 377–388, doi:10.1182/blood-2008-04-151712 (2009).

60 Zhang, L. et al. The RIG-I pathway is involved in peripheral T cell lymphopenia in patients with dermatomyositis. Arthritis Research & Therapy 21, 131, doi:10.1186/s13075-019-1905-z (2019).

61 Nabeshima, S. et al. A reduction in the number of peripheral CD28+CD8+T cells in the acute phase of influenza. Clinical & Experimental Immunology 128, 339–346, doi:10.1046/j.1365-2249.2002.01819.x (2002).

62 Nichols, J. E., Niles, J. A. & Roberts, N. J. Human Lymphocyte Apoptosis after Exposure to Influenza A Virus. Journal of Virology 75, 5921–5929, doi:10.1128/jvi.73.13.5921-5929.2001 (2001).

63 Gunzer, M. et al. Systemic administration of a TLR7 ligand leads to transient immune incompetence due to peripheral-blood leukocyte depletion. Blood 106, 2424–2432, doi:10.1182/blood-2005-01-0342 (2005).

64 Fathi, N. & Rezaei, N. Lymphopenia in COVID-19: Therapeutic opportunities. Cell Biology International 44, 1792–1797, doi:10.1002/cbin.11403 (2020).

65 Boonnak, K. et al. Lymphopenia Associated with Highly Virulent H5N1 Virus Infection Due to Plasmacytoid Dendritic Cell–Mediated Apoptosis of T Cells. The Journal of Immunology 192, 5906–5912, doi:10.4049/jimmunol.1302992 (2014).

66 Wei, C. J. et al. Next-generation influenza vaccines: opportunities and challenges. Nat Rev Drug Discov 19, 239–252, doi:10.1038/s41573-019-0056-x (2020).

