## Supplemental Information - Version 2 for "TLR7 and RIG-I dual-adjuvant loaded nanoparticles drive broadened and synergistic responses in dendritic cells *in vitro* and generate unique cellular immune responses in influenza vaccination"

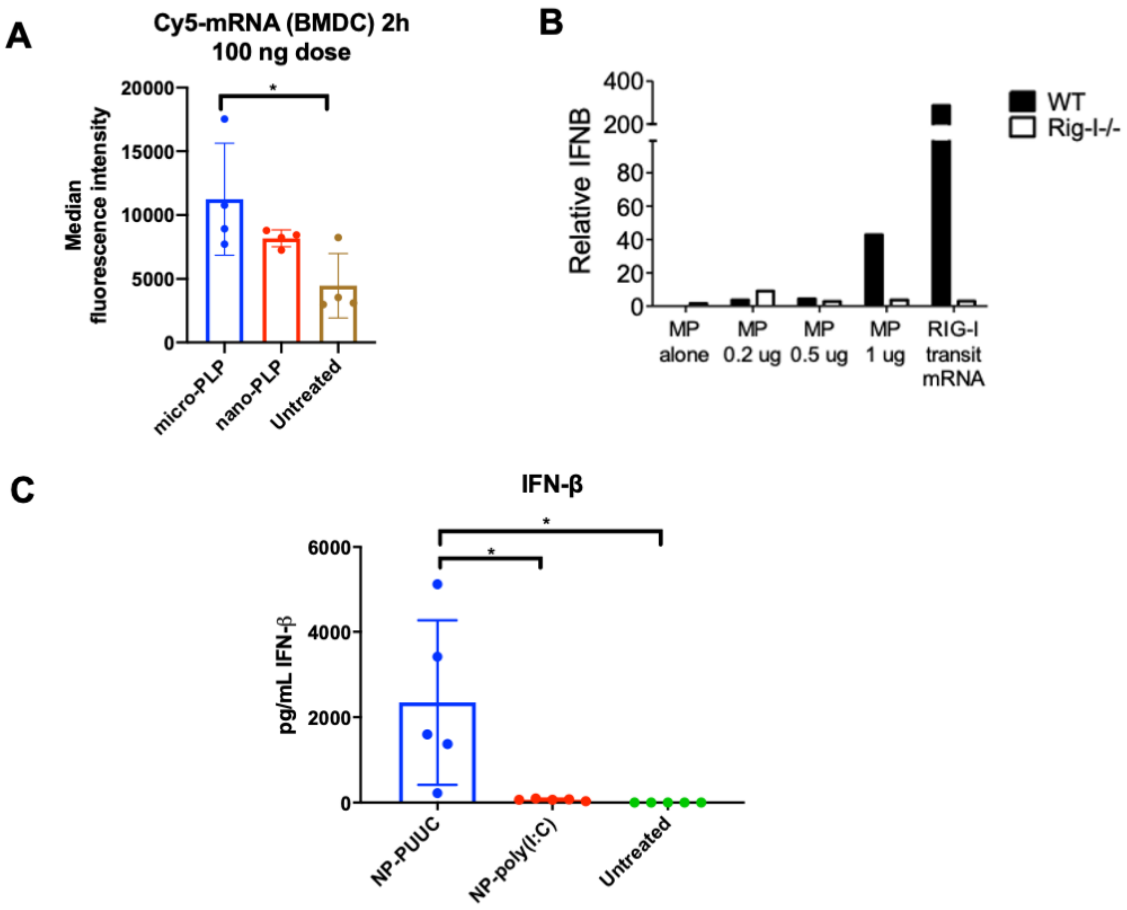


**SI Figure 1. Nano-PLPs enable delivery of functional RIG-I adjuvant (PUUC). A)** Median fluorescence intensity of mBMDCs treated with micro-PLPs or nano-PLPs with fluorescent luciferase mRNA. **B)** Comparison of wild-type and RIG-I$^{-/-}$ mBMDC activation after treatment with PUUC on micro-PLPs. **C)** Wild-type mBMDC activation (300,000 cells/well) with nanoparticles loaded with PUUC or poly(I:C). For both NP-PUUC and NP-poly(I:C), the loading level was 10 µg adjuvant/mg nanoparticles. Statistical differences were determined by one-way ANOVA followed by Tukey's test for multiple comparisons *P≤ 0.05.

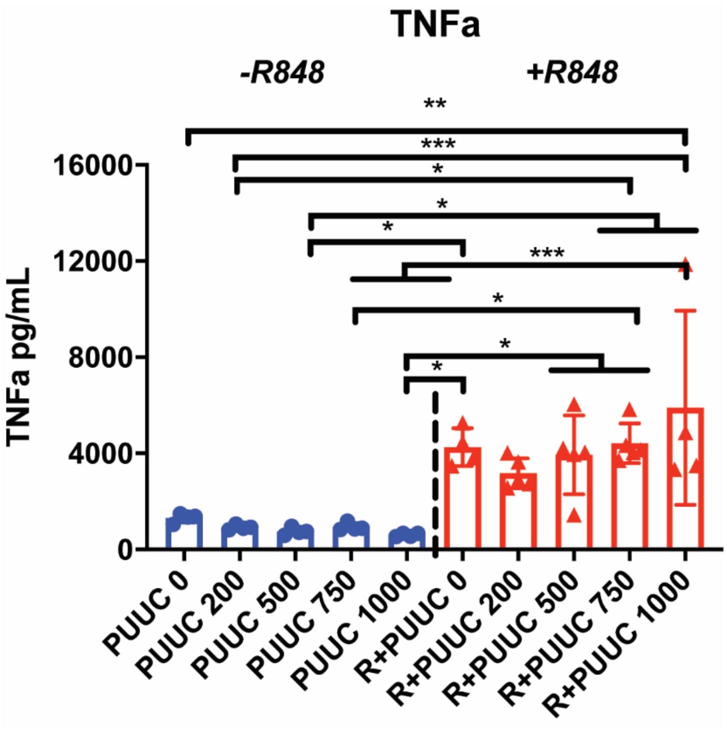


**SI Figure 2. TNF-α levels 24h after activation of pDCs.** CAL-1 human pDCs were treated with nanoparticles (86.5 µg/mL) loaded with R848 adjuvant (561 ng/mL) and PUUC adjuvant of doses ranging from 0-1000 ng/mL. TNFα supernatant concentrations were measured 24 hours after pDC activation. In all experiments, dual delivery was performed with a single nanoparticle system. Nanoparticle mass was fixed across all PUUC doses. Statistical significance was evaluated with one-way ANOVA followed by Tukey's test for multiple comparisons. $^*P \leq 0.05$, $^{**} P \leq 0.01$, $^{***}P \leq 0.001$.

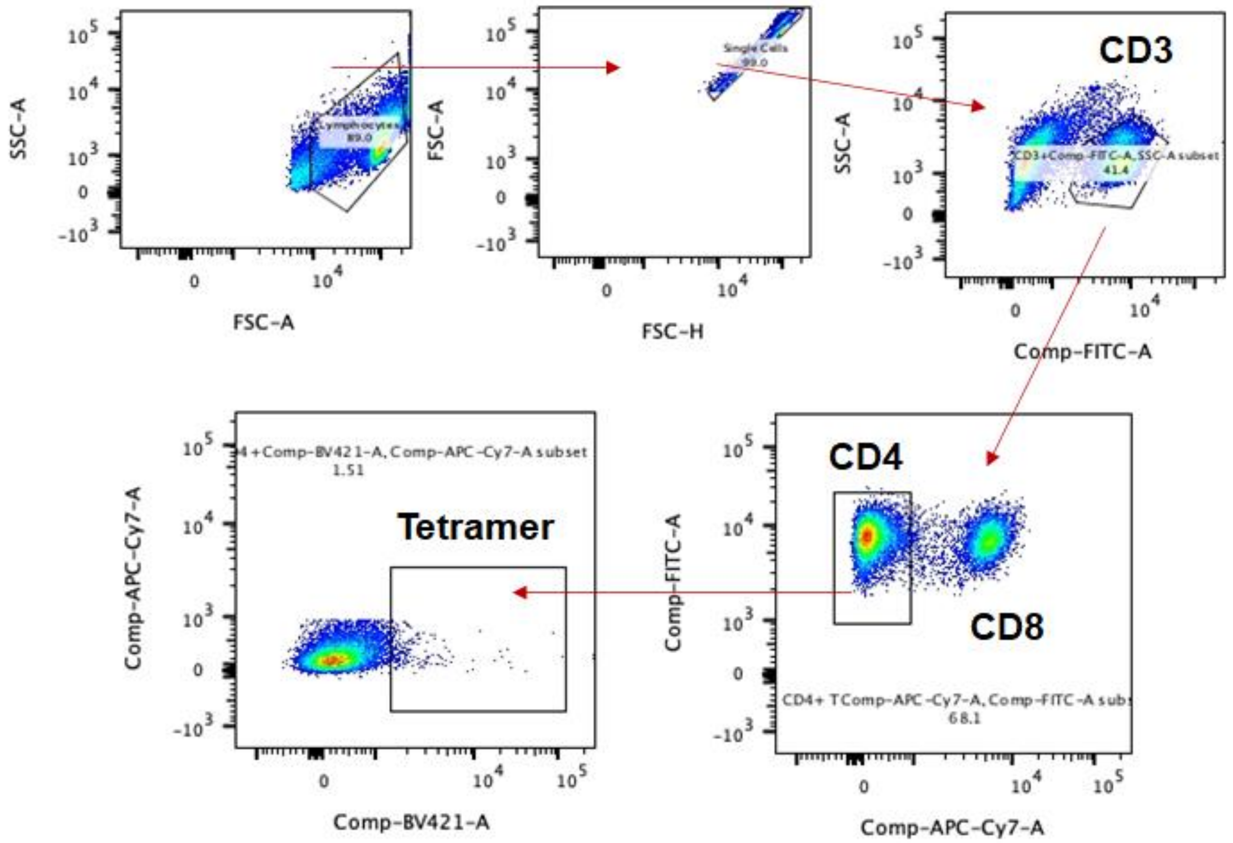


**SI Figure 3. Representative flow gating scheme to identify Class II tetramer positive CD4+ T-cells in the splenocytes.** The anti-CD4 antibody was not included in the flow cytometry panel, and therefore, the CD4+ T-cell was presumably gated on CD3+CD8- population.

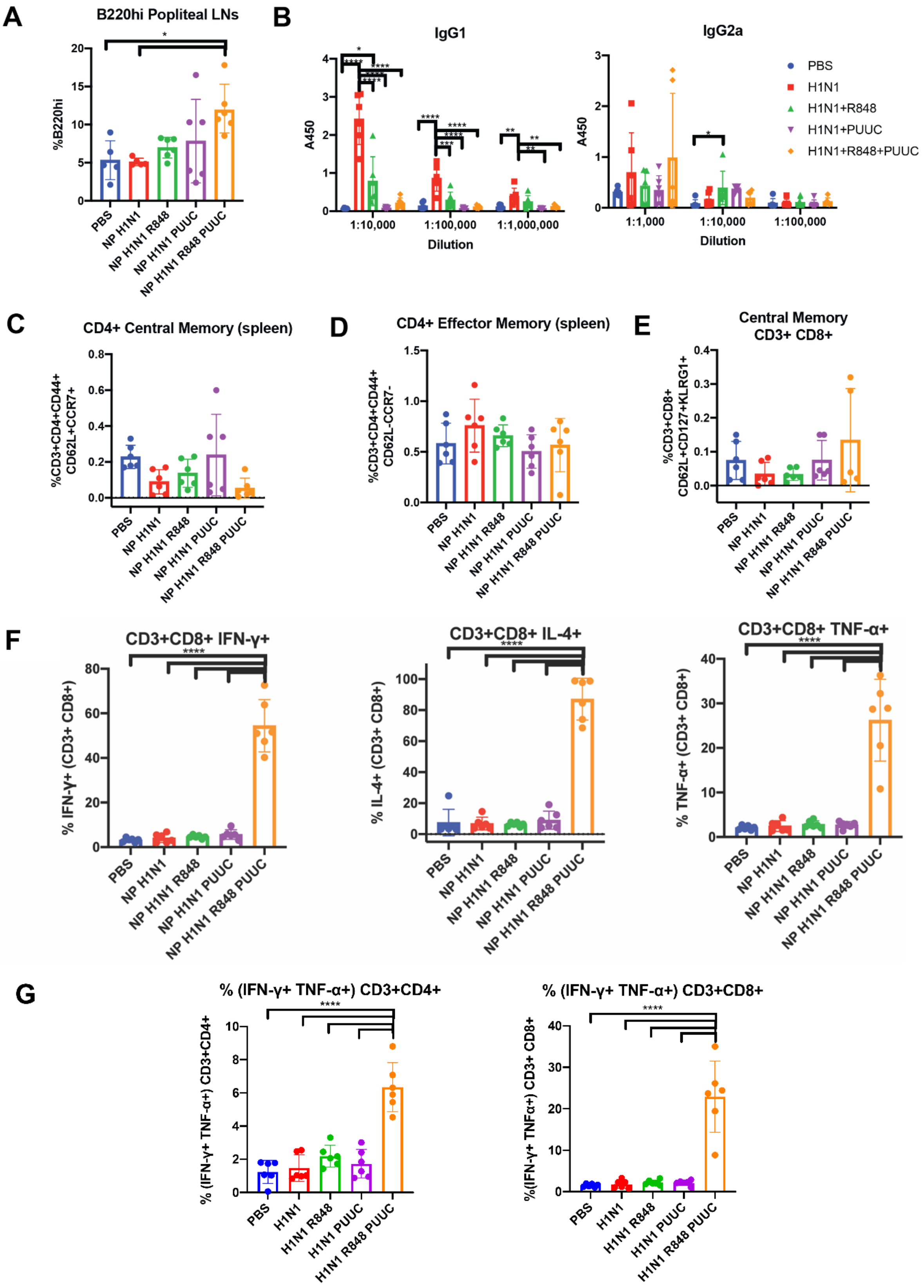
A
B220hi Popliteal LNs
%B220hi
PBS
NP H1N1
NP H1N1 R848
NP H1N1 PUUC
NP H1N1 R848 PUUC
B
IgG1
IgG2a
A450
1:10,000
1:100,000
1:1,000,000
1:1,000
1:10,000
1:100,000
Dilution
PBS
H1N1
H1N1+R848
H1N1+PUUC
H1N1+R848+PUUC
C
CD4+ Central Memory (spleen)
%CD3+CD4+CD44+ CD62L+CCR7+
D
CD4+ Effector Memory (spleen)
%CD3+CD4+CD44+ CD62L-CCR7-
E
Central Memory CD3+ CD8+
%CD3+CD8+ CD62L+CD127+KLRG1+
F
CD3+CD8+ IFN-γ+
% IFN-γ+ (CD3+ CD8+)
CD3+CD8+ IL-4+
% IL-4+ (CD3+ CD8+)
CD3+CD8+ TNF-α+
% TNF-α+ (CD3+ CD8+)
G
% (IFN-γ+ TNF-α+) CD3+CD4+
% (IFN-γ+ TNF-α+) CD3+CD4+
% (IFN-γ+ TNF-α+) CD3+CD8+
%(IFN-γ+ TNFα+) CD3+ CD8+
PBS
H1N1
H1N1 R848
H1N1 PUUC
H1N1 R848 PUUC

**SI Figure 4. Antibody and cell-mediated memory responses to NPs with HA, R848, and PUUC.** **A)** Percentage of popliteal lymphocytes with high expression of B220. **B)** IgG1 and IgG2a antibody titers were measured from serum. Populations of **C)** CD4+ central memory cells (CD3+ CD4+ CD44+ CD62L+ CCR7+), **D)** CD4+ effector memory cells (CD3+ CD4+ CD44+ CD62L- CCR7-), and **E)** CD8+ central memory cells (CD3+ CD8+ CD62L+ CD127+ KLRG1+) were measured in spleen. **F)** CD3+ CD8+ live splenocytes producing IFN-γ, IL-4, and TNF-α and **G)** polyfunctional CD4+ and CD8+ T-cells producing both IFN-γ and TNF-α after restimulation with H1N1 antigen for 6 hours**.** Error bars represent SD of the mean. Statistical significance was determined by one-way ANOVA followed by Tukey's test for multiple comparisons for normal datasets. $^{*}P \leq 0.05$, $^{**}P \leq 0.01$, $^{***}P \leq 0.001$, $^{****}P \leq 0.0001$.

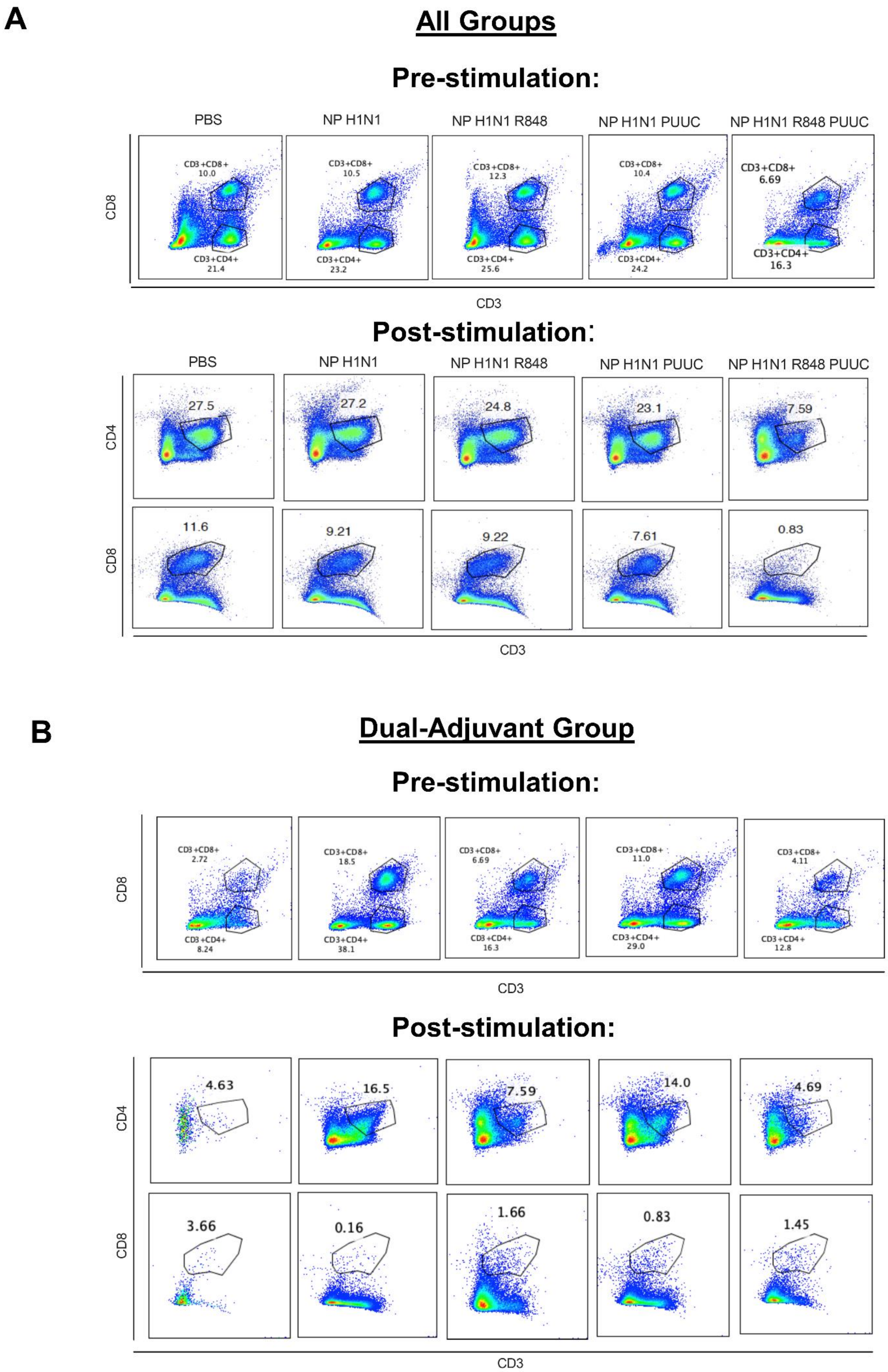

A
All Groups
Pre-stimulation:
PBS
NP H1N1
NP H1N1 R848
NP H1N1 PUUC
NP H1N1 R848 PUUC
CD8
CD3+CD8+
10.0
CD3+CD4+
21.4
CD3+CD8+
10.5
CD3+CD4+
23.2
CD3+CD8+
12.3
CD3+CD4+
25.6
CD3+CD8+
10.4
CD3+CD4+
24.2
CD3+CD8+
6.69
CD3+CD4+
16.3
CD3
Post-stimulation:
PBS
NP H1N1
NP H1N1 R848
NP H1N1 PUUC
NP H1N1 R848 PUUC
CD4
27.5
27.2
24.8
23.1
7.59
CD8
11.6
9.21
9.22
7.61
0.83
CD3
B
Dual-Adjuvant Group
Pre-stimulation:
CD8
CD3+CD8+
2.72
CD3+CD4+
8.24
CD3+CD8+
18.5
CD3+CD4+
38.1
CD3+CD8+
6.69
CD3+CD4+
16.3
CD3+CD8+
11.0
CD3+CD4+
29.0
CD3+CD8+
4.11
CD3+CD4+
12.8
CD3
Post-stimulation:
CD4
4.63
16.5
7.59
14.0
4.69
CD8
3.66
0.16
1.66
0.83
1.45
CD3

**SI Figure 5. Flow plots showing CD4+ and CD8+ T-cells in pre- and post-stimulated splenocytes with H1N1 antigen for single and dual adjuvant groups.** CD4+ and CD8+ T-cell population for **A)** all experimental groups and **B)** for dual-adjuvant (showing individual mice) treatment group before and after restimulation with H1N1 antigen for 6 hours. One replicate of the dual-adjuvant treatment group was omitted due to low staining. The anti-CD4 antibody was not included in the pre-stimulation flow cytometry panel, and therefore, the CD4 T-cell was presumably gated on CD3+CD8- population.
